# Neurovascular Uncoupling: Multimodal Imaging Delineates the Acute Effects of MDMA

**DOI:** 10.1101/2022.02.14.480365

**Authors:** Tudor M. Ionescu, Mario Amend, Tadashi Watabe, Jun Hatazawa, Andreas Maurer, Gerald Reischl, Bernd J. Pichler, Hans F. Wehrl, Kristina Herfert

## Abstract

Psychedelic compounds have attracted increasing interest in recent years due to their therapeutic potential for psychiatric disorders. Methylenedioxymethamphetamine (MDMA) is currently being investigated in clinical trials to treat post-traumatic stress disorder. To understand the acute effects of psychedelic drugs *in vivo*, functional MR imaging (fMRI) has been widely used in recent years. Notably, fMRI studies have shown that MDMA leads to inhibition of brain activity, challenging earlier hypotheses indicating mainly excitatory effects. However, interpretation of hemodynamic changes induced by psychedelics is challenging because of the potent vascular effects associated with this class of substances. Therefore, this study aimed to investigate the acute effects of MDMA using simultaneous positron emission tomography (PET)/fMRI in rats. For this purpose, hemodynamic changes measured by BOLD-fMRI were related to alterations in glucose utilization and serotonin transporter (SERT) occupancy, investigated using [^18^F]FDG functional PET (fPET) and [^11^C]DASB PET.

We demonstrate that MDMA induces global hemodynamic decreases accompanied by localized metabolic increases. Elevated metabolism was found primarily in limbic projection areas involved in emotion processing. Concurrent BOLD-fMRI decreases, also found in extracerebral areas, indicate that the BOLD-fMRI reductions observed in the brain are of vascular, non-neuronal origin. We further show that higher SERT occupancy strongly correlates with regional BOLD-fMRI reductions. Therefore, increased serotonin levels induced by SERT blockage may cause a neurovascular uncoupling.

Correct understanding of the *in vivo* mechanism of MDMA not only supports ongoing research but also warrants a reassessment of previous studies on neuronal effects of psychedelics relying on neurovascular coupling.

## 1. Introduction

In recent years psychedelic drugs, including lysergic acid diethylamide (LSD), psilocybin, and methylenedioxymethamphetamine (MDMA), have gained increasing attention due to their potential medical benefit for the treatment of psychiatric disorders [1, 2]. MDMA-assisted psychotherapy is currently in a phase three clinical trial to treat severe post-traumatic stress disorder (PTSD) with encouraging initial results [3]. Research in this area is also increasingly associated with the development of imaging techniques as quantitative biomarkers in addition to behavioral parameters [1]. To investigate the mechanisms of psychedelic drugs *in vivo*, magnetic resonance imaging (MRI) methods inferring neuronal activity through neurovascular coupling, such as blood oxygenation level-dependent functional MRI (BOLD-fMRI) and arterial spin labeling (ASL), have been widely used [4–8]. Interestingly, research performed over the last decade using the aforementioned methods has shown that psychedelic compounds such as MDMA [4] and psilocybin [6] inhibit brain activity, contradicting previous studies that indicated mainly excitatory effects [9–12].

However, the use of hemodynamic methods may be insufficient to understand the effects of psychedelics on neuronal activity. First, psychedelic drugs elicit their effects by strongly affecting one or more neurotransmitter systems [13]. Thus, it is crucial to evaluate hemodynamic changes in the context of changes in the respective neurotransmitter systems. Second, in addition to neuronal effects, increased neurotransmitter concentrations such as serotonin and dopamine elicited by psychedelic compounds can have potent vascular effects [14–16]. This aspect is particularly critical for methods based on neurovascular coupling, such as BOLD-fMRI and ASL. The emergence of hybrid positron emission tomography (PET)/MRI allows simultaneous assessment of brain function at multiple physiological levels [17, 18]. The combination of PET and pharmacological MRI (phMRI) [19] can provide high complementary measures of the effect of pharmacological compounds, especially for the evaluation of drug effects [20, 21]. In addition, recent developments in the administration of 2-[^18^F]fluoro-2-deoxyglucose ([^18^F]FDG) PET protocols via constant infusion [22] have paved the way towards functional PET (fPET) [22], a technique that allows imaging of changes in glucose metabolism at a temporal resolution of minutes [23]. fPET overcomes the lack of temporal resolution in previous studies, allows imaging of effects within a scan and provides a more robust indirect measure of neuronal activity compared to fMRI, because it is independent of changes in hemodynamics [22].

Here, we aimed to characterize the acute effects of MDMA using simultaneous PET/fMRI. First, we performed [^18^F]FDG fPET/fMRI scans to simultaneously determine hemodynamic and metabolic changes elicited by MDMA. In a second cohort, because the serotonin transporter (SERT) is one of the main targets of MDMA [24], we used the SERT PET ligand [^11^C]-3-amino-4-(2-dimethylaminomethylphenylsulfanyl)-benzonitrile ([^11^C]DASB) [25] to investigate whether SERT blockage by MDMA correlates with changes in BOLD-fMRI across the brain. The overall goal of this study was to exploit the potential of multimodal imaging for complementary detection and characterization of the acute *in vivo* effects of MDMA. By simultaneously acquiring data on hemodynamics, metabolism, and SERT occupancy, we aimed to test the two conflicting hypotheses that MDMA has either mainly inhibitory or excitatory effects on brain activity. A detailed description of the effects of MDMA will help to understand the mechanisms of the drug in the treatment of psychiatric disorders such as PTSD through MDMA-assisted psychotherapy, performed under acute MDMA.

## 2. Material and Methods

### Animals

Male Lewis rats (n = 29) were obtained from Charles River Laboratories (Sulzfeld, Germany) and divided into two groups: [^18^F]FDG fPET/fMRI scans were performed in 17 animals (361 ± 19 g), while [^11^C]DASB PET/fMRI were performed in 11 animals (365 ± 19 g). Nine fMRI datasets were excluded from the study due to motion during acquisition. One [^11^C]DASB PET and two [^18^F]FDG fPET datasets were excluded from the analysis because of paravenous tracer injections. The animals were kept at a room temperature of 22 °C and 40-60% humidity under a 12-hour light-dark cycle. The rats were fed with standard diet and received tap water *ad libitum*. Animals were fasted for six hours before the start of the experiments. All experiments were performed in accordance with the German Federal Regulations on the Use and Care of Laboratory Animals and approved by the Tübingen regional council.

Two additional sets of subjects scanned under the same [^18^F]FDG fPET/fMRI and [^11^C]DASB PET/fMRI protocols, but exposed to PBS instead of MDMA, are presented in the *supplementary information*.

### Simultaneous PET/fMRI experiments

Animals were placed in knock-out boxes, and anesthesia was induced with 3% isoflurane in regular air. Reflex tests were performed to determine sufficient sedation. For subsequent preparation procedures, the isoflurane concentration was reduced to 2%. After the weight of the animals was determined, a catheter with a 30-G needle was positioned in a tail vein to administer the radioactive tracer. Another catheter was placed into the other tail vein for MDMA administration. Animals were then placed on a dedicated temperature-controlled small animal bed (Medres, Cologne, Germany). The temperature was monitored and maintained at 36.5°C with a rectal probe, and respiratory rates were monitored using a breathing pad. Rats were then placed in the scanner, and the isoflurane concentration was decreased to 1.3% throughout the scan.

Scans were performed using a 7-T small-animal MRI scanner (ClinScan, Bruker Biospin, Ettlingen, Germany) using a 72-mm-diameter linearly polarized RF coil (Bruker) for transmission and a four-channel rat brain coil (Bruker) for reception. First, localizer scans were performed to position the brains in the center of the field of view. Local magnetic field maps were then generated to optimize local field homogeneity. Subsequently, T2-weighted MRI sequences (TR: 1800 ms, TE: 67.11 ms, FOV: 40 × 32 × 32 mm^3^, image size: 160 × 128 × 128 px, Rare factor: 28, averages: 1) were performed to obtain anatomical references. Finally, fMRI imaging was performed using T2*-weighted gradient-echo EPI sequences (TR: 2000ms, TE: 18ms, 0.25 mm isotropic resolution, FoV 25 × 23 mm^2^, image size: 92 × 85 × 20 px, slice thickness: 0.8 mm, 20 slices).

Simultaneously, PET scans were acquired using a PET insert developed in collaboration with Bruker, which is the second generation of an insert previously developed in-house [18]. Protocols for tracer synthesis, injected radioactivities and molar activities are given in *supplementary information*. PET and fMRI acquisitions were started simultaneously 30 seconds before tracer injection and performed over 100 minutes after tracer injection. PET data were saved as list-mode files for later reconstruction into dynamic scans of 100 1-minute frames using an ordered-subsets expectation maximization 2D (OSEM-2D) algorithm.

A pharmacological MDMA challenge of 3.2 mg/kg was administered 40 minutes after tracer injection over 30 seconds.

### Data preprocessing

Statistical Parametric Mapping 12 (SPM 12, Wellcome Trust Centre for Neuroimaging, University College London, London, United Kingdom) via Matlab (The MathWorks, Natick, MA, USA) and Analysis of Functional NeuroImages (AFNI, National Institute of Mental Health (NIMH), Bethesda, Maryland, USA) were used for data preprocessing as previously reported [26]. An extensive description of all preprocessing steps performed can be found in the *supplementary information*.

To analyze the effect of non-neuronal signal changes on the brain BOLD-fMRI measures, we extracted the average signal from all voxels in areas outside the brain using masks created in AFNI. Average time courses were extracted from all datasets after preprocessing using the MarsBaR toolbox [27] and regions of interest defined by the Schiffer brain atlas [28]. For a complete list of the regions, please refer to *supplementary information*.

### SPM data analysis

The general linear model (GLM) available in SPM was applied to determine voxels with significantly altered fMRI and PET signals after MDMA exposure. For all datasets, the baseline was defined as the period between 30 and 40 minutes after scan start, after tracer equilibrium had been reached between the high target and the reference regions. The [^18^F]FDG fPET data were normalized based on cerebellar uptake, as previously recommended [29], whereas for [^11^C]DASB PET, the cerebellar gray matter was chosen as the reference region [30].

For PET data analysis, mean images of the dynamic 1-minute frames were generated over seven measurement periods: between 30 and 40 minutes following tracer injection as baseline, and for each of the subsequent 10-minute periods after MDMA exposure (40-50 minutes, 50-60 minutes, 60-70 minutes, 70-80 minutes, 80-90 minutes and 90-100 minutes after scan start). Paired t-maps were then calculated between the baseline and each block after MDMA for [^18^F]FDG using voxel-wise normalized uptake maps and for [^11^C]DASB using voxel-wise DVR-1 maps. For more details please refer to *supplementary information*.

For fMRI, a first-level analysis was applied to the individual scans using the pseudo-block approach reported for phMRI [31, 32]. To allow a direct comparison with the PET readout, the same six 10-minute blocks between MDMA challenge and the end of the scans (40-50 minutes, 50-60 minutes, 60-70 minutes, 70-80 minutes, 80-90 minutes, and 90-100 minutes after scan start) were compared with the baseline 10-minute block before MDMA injection. After estimating the GLM parameters, statistical parametric maps were generated by interrogating the data using contrast vectors between each block after MDMA and baseline. The generated maps were then used to delineate group-level effects through a second-level analysis. Initially, binary masks were used to perform the analyses only within the brain. The binary masks were removed for later analysis of extracerebral hemodynamic alterations.

All group-level t-maps were subjected to voxel-wise signal quantification to determine the regional contributions of brain regions selected according to the Schiffer brain atlas [28]. The average t-scores of all voxels comprising each region were calculated for each period and modality to compare the respective spatial patterns of MDMA effects on hemodynamics, glucose metabolism, and SERT occupancy.

## 3. Results

### Metabolic increases accompany hemodynamic reductions after MDMA

First, we investigated the relationship between acute hemodynamic and metabolic changes following acute MDMA using a simultaneous [^18^F]FDG fPET/fMRI protocol (Figure 1).

**Figure 1:**
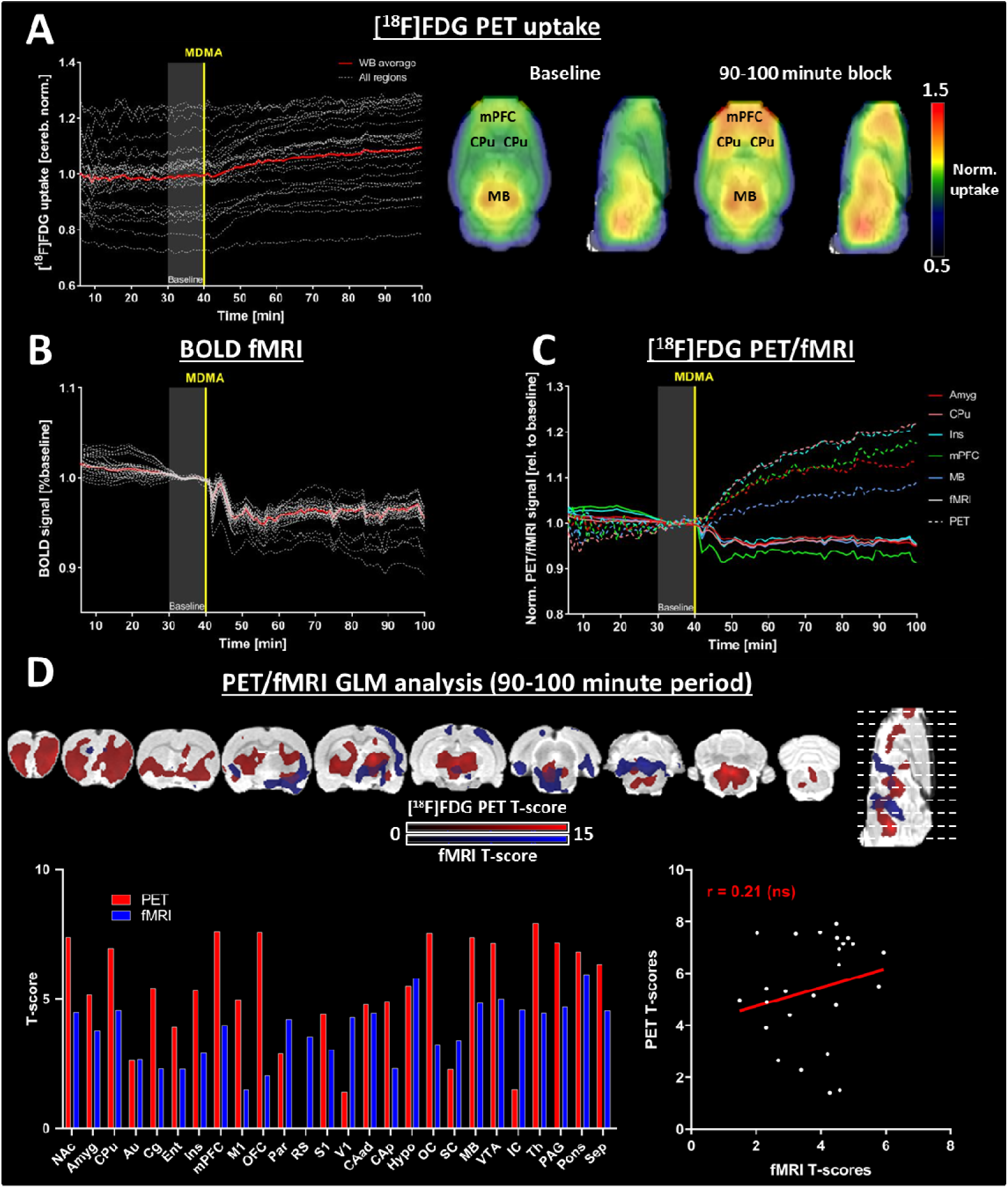
Region-wise and voxel-wise evaluation of [^18^F]FDG fPET and BOLD-fMRI signals changes. (A) TACs for all regions and whole-brain average. Voxel-wise normalized uptake maps indicate [^18^F]FDG uptake at baseline (30 to 40 minutes after scan start), as well as 50-60 minutes after MDMA administration. (B) Regional BOLD-fMRI signals normalized to respective baselines (average regional BOLD-fMRI signals in the period 30-40 minutes after scan start). (C) Both signals were normalized to the baseline period over the last ten minutes before MDMA administration for a common frame of reference. Continuous lines indicate the BOLD-fMRI signals, interrupted lines the [^18^F]FDG fPET signals in Amyg, CPu, Ins, mPFC, and MB. (D) Voxel-wise analysis of both signals in the period 50-60 minutes after MDMA challenge. The voxelwise maps are presented at p < 0.05 (FWE-corrected at voxel level) for PET and p < 0.001 at voxel level with p < 0.05 FWE correction at cluster level for fMRI (n = 15 for fPET, n = 9 for fMRI). The bar diagram indicates average t-scores for each region and both modalities. The average regional t-scores are plotted in a scatter diagram to evaluate the spatial correlation of both readouts. Abbreviations: GLM = general linear model, ns = not significant. For abbreviations of the different regions, please refer to *Supplementary Information*.

The normalized [^18^F]FDG fPET TACs of all regions and the average whole-brain TAC are shown in Figure 1A. Visual assessment of TACs and voxel-wise uptake maps indicated an increase in metabolism in the midbrain and subcortical areas such as the striatum and frontal cortical areas, whereas more minor or no changes occurred in posterior cortical regions. Notably, we found a simultaneous decrease in hemodynamics, as indicated by BOLD-fMRI (Figure 1B). The whole-brain averaged BOLD-fMRI signal was reduced by 4.5% fifteen minutes after the challenge. Importantly, the data revealed that the decreases were of global nature and occurred in all regions investigated.

A temporal comparison of the [^18^F]FDG fPET and BOLD signal changes relative to baseline is shown in Figure 1C. The highest metabolic increases occurred in frontal areas, including the CPu (22% increase 60 minutes post-challenge), Ins (21% increase 60 minutes post-challenge), mPFC (18% increase 58 minutes post-challenge), and Amyg (13.5% increase 58 minutes post-challenge). Temporally, increases in all regions (>1%) were observed within 5 minutes of challenge.

The voxel-wise GLM analyses presented in Figure 1D revealed that the metabolic rate increased across several subcortical areas and in frontal cortical areas in the period between 50 and 60 minutes after the challenge. The mPFC and OFC (t = 7.6 for both), along with MB (t = 7.4), Th (t = 7.9) and NAc (t = 7.4) exhibited the most significant [^18^F]FDG increases. The most significant BOLD-fMRI decreases occurred in posterior areas such as the MB (t = 4.8), VTA (t = 5.0), Hypo (t = 5.8), and Pons (t = 5.9). The t-scores of metabolic increases and BOLD-fMRI decreases did not correlate significantly (r = 0.21), underlining a disturbed relationship between metabolism and hemodynamics.

### SERT occupancy changes induced by MDMA correlate with BOLD decreases

To further elucidate the molecular underpinnings of the observed hemodynamic decreases, we evaluated BOLD-fMRI changes concurrently with alterations in SERT availability using [^11^C]DASB PET/fMRI in a second cohort (Figure 2).

**Figure 2:**
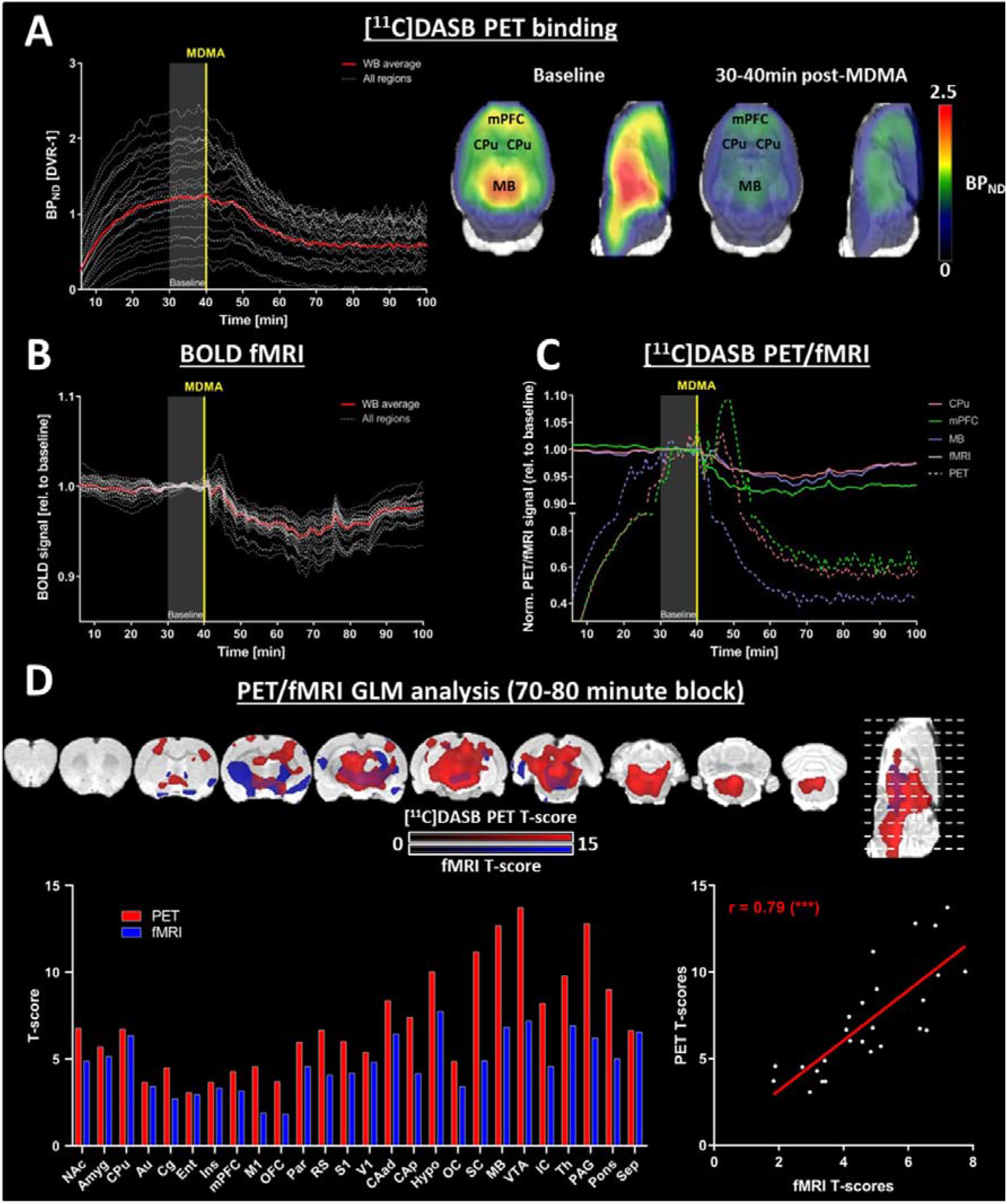
Region-wise and voxel-wise evaluation of [^11^C]DASB PET and BOLD-fMRI signal changes. **(A)** Dynamic binding potentials for all regions and whole-brain average. Voxel-wise binding potential maps indicate [^11^C]DASB binding at baseline (30 to 40 minutes after scan start), as well as 70-80 minutes after scan start (30-40 minutes post-challenge). **(B)** Regional BOLD-fMRI signals normalized to respective baselines (average regional BOLD-fMRI signal in the period 30-40 minutes after scan start). **(C)** Temporal comparison of PET and BOLD signal changes (normalized to the baseline) in CPu, mPFC, and MB. **(D)** Voxel-wise analysis of both signals in the period 30-40 minutes after MDMA challenge. The voxelwise maps are presented at p < 0.05 (FWE-corrected at voxel level) for PET and p < 0.001 at voxel level with p < 0.05 FWE correction at cluster level for fMRI (n = 11). The bar diagram indicates average t-scores for each region and both modalities. The average regional t-scores are plotted in a scatter diagram to evaluate spatial correlation of both readouts (*** indicates significance at p < 0.001). Abbreviation: GLM = general linear model; for abbreviations of the different regions please refer to *Supplementary Information*.

The DVR-1 of [^11^C]DASB reached equilibrium 30 minutes after injection (Figure 2A). After pharmacological challenge with MDMA, binding values in all regions decreased either immediately (1-2 minutes after the challenge) in areas with high binding values (BP_ND_ > 1.8) or with a delay after approximately 10 minutes in regions with lower [^11^C]DASB binding values. At 30 minutes post-challenge, the binding values remained stable until the end of the scan period. Voxel-wise binding values reflected the most significant decreases of [^11^C]DASB binding in regions with high SERT availability. Similarly to the BOLD-fMRI dataset acquired in the [^18^F]FDG fPET/fMRI cohort, all regional BOLD signals decreased within 6 minutes of MDMA challenge (Figure 2B). Fifteen minutes after the MDMA challenge, the whole-brain average BOLD signal was decreased by 4% compared to baseline.

A temporal comparison of the changes in [^11^C]DASB binding and the BOLD-fMRI responses for three exemplary regions (Figure 2C) revealed that regions with higher baseline SERT availability showed a faster response than regions with lower baseline SERT availability. For example, [^11^C]DASB binding in the MB, a region with high baseline [^11^C]DASB binding (BP_ND_ = 2.1), decreased one minute after challenge reaching 59% of its baseline value at 40 minutes after the challenge. In contrast, [^11^C]DASB binding in the CPu (BP_ND_ = 1.6) and mPFC (BP_ND_ = 1.6) remained stable or increased shortly after the MDMA challenge. After approximately 10 minutes, [^11^C]DASB binding decreased in all regions until it reached equilibrium at 30-40 minutes after the challenge (39% decrease for mPFC, 44% decrease for CPu 40 minutes post-challenge compared to baseline). Compared to the temporally differentiated [^11^C]DASB binding reductions delineated above, decreases in BOLD occurred homogeneously. The highest regional BOLD decreases occurred within the first 30 minutes after challenge. Reductions between 1% and 2% could be observed for mPFC, CPu, and MB as early as 2 minutes after challenge. The decreases peaked within 30 minutes after MDMA injection (~8.5% in mPFC, ~5.5% in CPu, and ~6.5% in MB).

Figure 2D shows GLM analyses for both readouts between baseline and 70-80 minutes after the start of the scan, when changes reached a plateau for both methods. [^11^C]DASB decreases were highest in the VTA (t = 13.7), PAG (t = 12.8) and MB (t = 12.7), regions with high baseline [^11^C]DASB binding values. The largest regional t-scores for BOLD-fMRI occurred in the Hypo (t = 7.8), VTA (t = 7.2) and Th (t = 6.9). Remarkably, regional t-scores of both readouts correlated strongly (r = 0.79, p < 0.001), indicating a strong relationship between SERT occupancy and BOLD signal decreases.

### Hemodynamic reductions also occur in non-neuronal tissues

The smaller spatial extent of hemodynamic changes compared with the metabolic and SERT occupancy changes presented for the two cohorts above may be due to the smaller magnitudes of the BOLD decreases. To further clarify this aspect, we merged the fMRI scans from both cohorts (Figure 3). We also extracted the BOLD signals from extracerebral areas to investigate whether the BOLD decreases are specific to neuronal tissue. Finally, we compared the temporal characteristics of hemodynamic, metabolic, and SERT occupancy changes.

**Figure 3:**
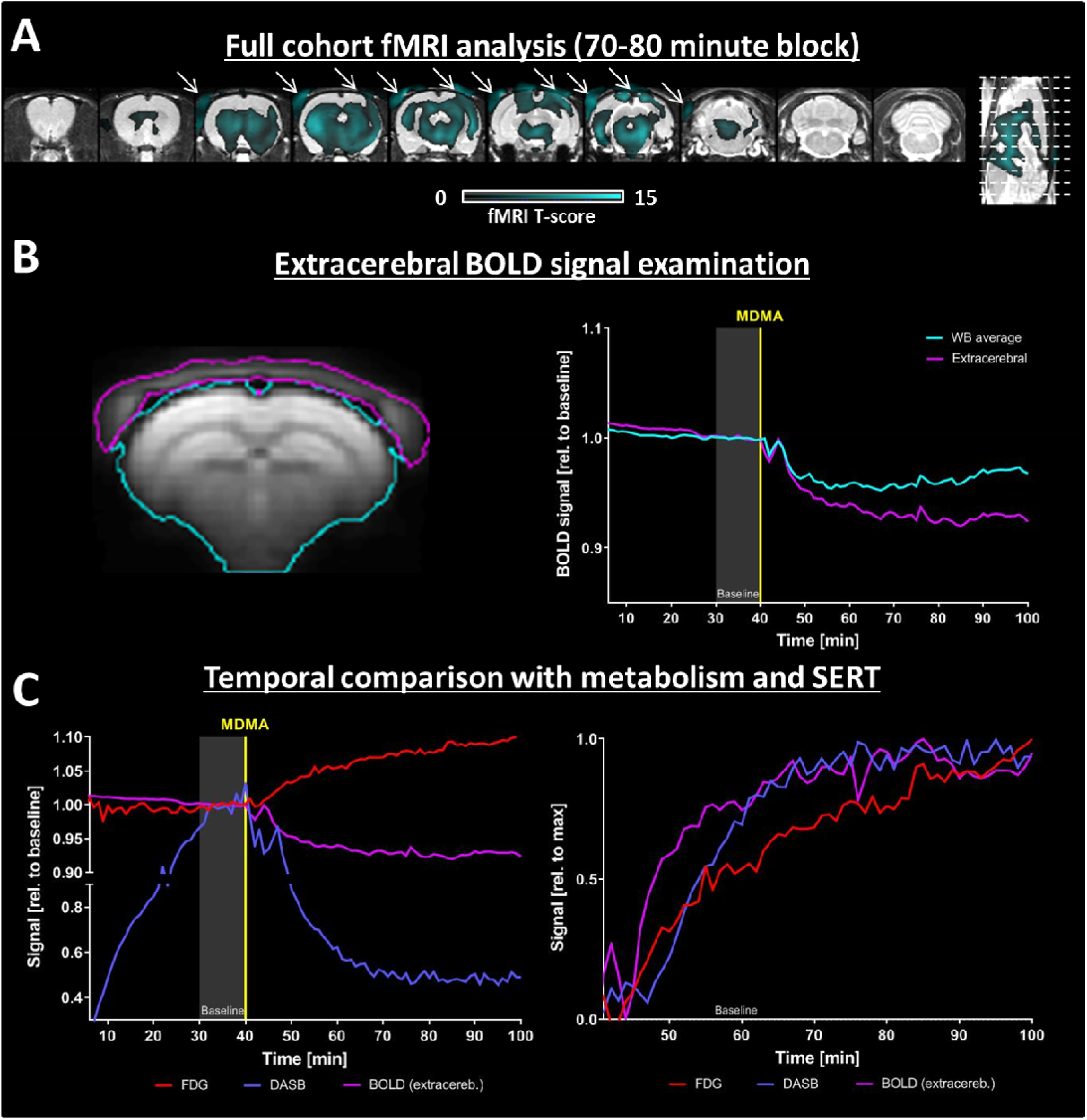
Examination of BOLD signal changes in extracerebral areas. **(A)** GLM analysis of BOLD decreases after merging BOLD-fMRI datasets acquired in both [^18^F]FDG and [^11^C]DASB cohorts (n=20). Data are shown at p < 0.05, FWE-corrected at the voxel level. Arrows indicate extracerebral decreases. **(B)** BOLD signals were extracted from the extracerebellar regions (indicated in cyan), averaged over the cohort, and plotted along with the group-average whole-brain signal. **(C)** Left: Temporal alterations of the three signals in relation to the baseline. Right: Temporal changes of the readouts after MDMA challenge relative to their respective maximum changes (0 = no change, 1 = maximum change) are shown in order to depict the temporal evolution of the alterations.

Because of the global nature of the observed hemodynamic decreases, we also investigated changes in BOLD signal without masking extracerebral areas (Figure 3A). The analysis indicates the decrease in BOLD-fMRI is widespread and comparable in magnitude to the decrease in [^11^C]DASB PET binding and, intriguingly, also occurs in non-neuronal areas. Thus, we extracted BOLD signals from extracerebral areas consisting predominantly of tissue surrounding the skull (Figure 3B) to determine the temporal characteristics of the observed decreases. Remarkably, the extracerebral BOLD signal decreased coherently with the cerebral BOLD signal, both reaching maximum reductions 27 minutes after the challenge (4.8% for cerebral BOLD signal and 7.4% for extracerebral BOLD signal).

The temporal profiles of the changes in hemodynamics, metabolism, and SERT occupancy are shown in **Error! Reference source not found**. C and D, respectively. The temporal characteristics of hemodynamic and SERT availability decreases were highly comparable, both reaching 90% of their maximum changes 30 minutes after the challenge. In contrast to the other two datasets, [^18^F]FDG increased linearly, reaching 90% of the maximum change only 45 minutes after the challenge, with the whole-brain average [^18^F]FDG TAC continuing to increase until the end of the scan. This finding suggests that metabolic increase may persist longer than one hour after the MDMA challenge included in the scans.

## 4. Discussion

In this study, we investigated the acute effects of the psychedelic drug MDMA using simultaneous PET/fMRI. For this purpose, we examined the relationships between changes in glucose metabolism, vascular responses, and SERT availability in different brain regions. We detected concurrent localized increases in glucose consumption and hemodynamic decreases of global nature, both occurring within minutes after MDMA exposure. Furthermore, as expected, we identified decreased [^11^C]DASB specific binding to the SERT following the challenge. Remarkably, regional SERT changes correlated strongly with decreased hemodynamics, whereas increased glucose metabolism measured by [^18^F]FDG fPET showed no correlation with BOLD reductions. Finally, we show that hemodynamic decreases concurrently take place in non-neuronal extracerebral tissues. Our data suggest that increased neuronal activity is accompanied by neurovascular uncoupling, possibly mediated through the vascular effects of serotonin following SERT blockage.

### Simultaneous uncoupling between metabolism and hemodynamics

The acute *in vivo* effects of psychedelic compounds, and MDMA in particular, have not been fully elucidated in terms of neuronal activation or inhibition, with two hypotheses being postulated. *In vivo*, studies measuring glucose utilization have shown mixed results, yet mainly increased metabolism following psychedelic challenges [9, 10, 12, 33], suggesting neuronal activation. However, more recent work with ASL challenged this hypothesis and argued that brain activity is decreased under MDMA [4]. Interestingly, the authors did not find any hemodynamic increase following MDMA across the entire brain. Our study confirms that this finding also holds when applying an acute MDMA challenge in rodents. Furthermore, the authors speculated that MDMA exerts inhibitory effects directly through 5-HT_1A_ receptors, which are known to induce hyperpolarization and decrease firing rates [4, 34]. The same group found similar hemodynamic decreases under acute psilocybin [6], postulated by the authors to be driven by inhibitory effects of 5-HT_2A_ receptors [35]. On a side note, the decreases in ASL and BOLD previously observed for MDMA and psilocybin in humans were also asymmetric [4, 36], with a focus on the right cerebral hemisphere. The same pattern can be observed in our study, supporting the translatability of our readout. Notably, the authors argued that possible discrepancies with previous work indicating increased metabolism using [^18^F]FDG PET [12] might be because [^18^F]FDG PET operates on a much longer time scale than fMRI [6]. Therefore, the authors claimed that the increases in earlier [^18^F]FDG PET studies may represent a rebound in glucose metabolism after the actual acute inhibitory effects captured by fMRI [6]. We agree with the above statement that earlier [^18^F]FDG PET or *ex vivo* studies [9, 10, 12, 33] measuring cerebral glucose utilization lacked temporal specificity because of the paradigms used, which complicates the interpretation of results for acute effects. Earlier work on blood flow effects is similarly inconclusive and partly confirmes the general decreases induced by MDMA seen with ASL [4, 9]; however, CBF measurements using [^15^O]H_2_O PET indicated mixed regional increases and decreases [37]. A further reason for the partially conflicting results of the above studies is that the conclusions were drawn between different, relatively small cohorts having received either placebo or MDMA.

In our study, the above limitations of previous studies were overcome. Recently developed PET protocols have enabled the detection of metabolic effects at high temporal resolutions of only a few minutes [22, 23]. Our similarly designed study allowed temporal assessment of the acute effects of two pairs of readouts: metabolic versus hemodynamic response and SERT occupancy versus hemodynamics. The effects were delineated (1) immediately after the challenge at 1-minute intervals, (2) simultaneously for both pairs of measurements, and (3) in the same animals, thus avoiding the comparison of two different groups of subjects. Therefore, we demonstrate that the uncoupling between flow and metabolism suggested in previous work [9, 14, 33] does occur in the same subjects and almost simultaneously.

### Origin of peripheral and cerebral hemodynamic effects

We show that non-neuronal effects dominate hemodynamic changes induced by MDMA. In particular, our data shed light on two separate phenomena. First, temporal coherence between hemodynamic reductions in cerebral and extracerebral areas suggests that vascular effects occur in the periphery. The 5-HT_2A_, which is postulated, along with the 5-HT1B receptor, to mediate vasoconstrictive effects [38, 39], is one of the main targets of MDMA [5, 36]. However, because MDMA has a much stronger affinity to the 5-HT_2A_ receptor than to the 5-HT_1B_ receptor [5], direct agonist action of MDMA at the 5-HT_2A_ in peripheral blood vessels likely results in vasoconstriction [38]. This is supported by work showing that the use of 5-HT_2A_ antagonists following the elevation of serotonin levels reduces the serotonin-induced vasoconstriction in the carotid artery, the main vessel that provides blood supply to the brain [40]. Furthermore, compounds with high selectivity for 5-HT_2A_ have been shown to increase blood pressure in carotid arteries, indicating vasoconstrictive effects [41].

In addition to the peripheral effects of MDMA, the reductions observed in the brain are likely also of direct serotonergic nature, consistent with the previously reported effects of serotonin on brain microvasculature [15, 16]. Early work has shown that manipulation of the raphé nuclei directly affects brain microcirculation by inducing vasoconstriction [14]. Interestingly, although both MDMA and psilocybin act potently on the serotonergic system, psilocybin has a very low affinity to the SERT, unlike MDMA [42, 43]. Based on this fact, it is interesting that both drugs appear to induce similar spatial patterns of hemodynamic decreases [4, 6]. Since of all serotonergic receptors, psilocybin has the highest affinity to the 5-HT_2A_ receptor [43], which has been shown to induce vasoconstriction peripherally, it is reasonable to assume that this is similar in the brain. In contrast to the peripheral effects of MDMA, which can be attributed solely to the direct effects of the drug, the observed decreases in the brain may additionally be triggered by increased synaptic serotonin levels following SERT blockage. This finding is supported by the very high correlation between SERT blockage and hemodynamic decreases in BOLD-fMRI, suggesting that direct effect of endogenous serotonin may additionally modulate hemodynamic decreases in the brain. In the future, further insight may be gained by combining a psychedelic challenge with antagonists to 5-HT_2A_ and other serotonin receptors to elucidate their respective involvements in the observed hemodynamic decreases.

Our results warrant a re-evaluation of the hypothesis that the observed hemodynamic reductions by psychedelic drugs are driven by neuronal activity, which is also supported and discussed in later publications [44–47]. In general, additional caution is necessary when interpreting findings relying on neurovascular coupling, such as BOLD-fMRI functional connectivity studies, under pharmacological challenges [4, 5, 36].

### Spatial and temporal properties of increased metabolism

We demonstrate that MDMA triggers increased glucose consumption, likely due to neuronal activation, in different regions by [^18^F]FDG fPET measurements. Changes in [^18^F]FDG fPET have been shown to reliably reflect changes in neuronal activity while being independent of hemodynamic changes [22]. Our data revealed a substantial increase in metabolic activity that started a few minutes after MDMA injection and extended from the raphé nuclei to anterior subcortical and cortical projection areas. Interestingly, although many increases in glucose consumption overlapped with SERT blockage, the metabolic increases were more weighted toward projection areas than the reductions seen in [^11^C]DASB binding. First, this finding is in line with the hypothesis that the majority of glucose is consumed postsynaptically [48, 49]. Second, the areas showing increased metabolism are consistent at a functional level with the majority of previously reported behavioral effects of MDMA. The signals observed in the nucleus accumbens, amygdala, and insula compare well with salience changes known from imaging and behavioral studies [4, 50]. In particular, the nucleus accumbens, as the main reward processing hub, is involved in responses to numerous drugs [51]. In addition, the amygdala, insula, and orbitofrontal cortex are strongly involved in emotional processes [52], so their increased activities could explain the enhanced emotionality and empathy reported as effects of the drug [53]. Activity in the olfactory cortex and olfactory bulb could indicate increased food-seeking or sexual arousal [53, 54]. Enhanced metabolic activity in sensory cortices is in concordance with heightened sensations elicited by MDMA [53]. At the molecular level, the aforementioned 5-HT_2A_ receptor exhibits a strong anterior-posterior gradient in the cortex, with strong expression in frontal areas and less expression in posterior areas of the cortex [55], consistent with the activations indicated by our [^18^F]FDG PET data, predominantly in frontal cortical areas. Moreover, the 5-HT_2A_ receptor has been shown to be responsible for serotonergic activation in projection areas such as the prefrontal cortex [56, 57]. However, the exact receptors involved in neuronal activation indicated by fPET can only be speculated because both MDMA and endogenous serotonin may play a role, both of which can bind to a wide array of monoaminergic receptors.

Temporally, changes in [^11^C]DASB and extracerebral BOLD predominated in the first 30 minutes after the challenge, both reaching over 90% of their respective maximal changes at this time point. Therefore, the two readouts likely reflect the direct molecular and vascular effects of MDMA discussed above. This finding is consistent with previous work indicating peak serotonin release 20 minutes after intraperitoneal MDMA injection [13]. In contrast, increases in [^18^F]FDG fPET, likely reflecting increased neuronal activity, did not reach a plateau 60 minutes after the challenge at the end of the experiment, suggesting prolonged activation after the peak of direct MDMA action.

### Further considerations

Other factors may play a role in the findings. First, MDMA and endogenous serotonin bind to other receptors highly expressed in the brain, such as the 5-HT1 receptor family [5, 58], which could also be involved in the observed effects. Next, the neuronal activations indicated by fPET could be caused by the effects of MDMA on other neurotransmitters, primarily norepinephrine and dopamine [13, 59]. Dopamine, similarly to serotonin, has been shown to cause vasoconstriction when released at high concentrations [60]. However, the strong correlation between SERT occupancy and hemodynamic changes implies that serotonin is more actively modulating brain microcirculation after acute MDMA application than other monoamines, as previously postulated [9, 14, 33]. To understand how exactly regional hemodynamic and metabolic changes occur, distributions of all drug targets are generally required. PET/MRI imaging offers the opportunity to elucidate the effects of psychedelic drugs by combining data on transporters and receptors with hemodynamics measured by BOLD-fMRI. Recent work has demonstrated how insight into different receptor distributions can be used to understand changes in BOLD-fMRI data [5]. Such analytical approaches are of interest to gain a deeper understanding of drug mechanisms of action. A more thorough discussion on the advantages and limitations of multimodal imaging and small-animal imaging can be found in *supplementary information*.

## 5. Conclusion

The present study pioneers and demonstrates the tremendous potential of multimodal imaging in psychedelic drug research. We demonstrate the neurovascular uncoupling induced by an acute MDMA challenge, characterized by increased neuronal activity in monoaminergic projection areas accompanied by vascular depression caused, at least in part, by serotonin release. Our results provide tremendous insight into the mechanism of action of MDMA and pave the way for the application of hybrid PET/fMRI in psychedelic drug research.

## CRediT authorship contribution statement

**Tudor M. Ionescu:** Conceptualization, Methodology, Software, Validation, Formal analysis, Data curation, Writing – original draft, Visualization. **Mario Amend:** Conceptualization, Methodology, Investigation, Writing – review & editing, Supervision. **Tadashi Watabe:** Methodology, Investigation, Writing – review & editing. **Jun Hatazawa:** Writing – review & editing. **Andreas Maurer:** Writing – review & editing. **Gerald Reischl:** Writing – review & editing. **Bernd J Pichler:** Writing – review & editing, Funding acquisition. **Hans F. Wehrl:** Conceptualization, Methodology, Writing – review & editing, Funding acquisition, Visualization. **Kristina Herfert:** Methodology, Writing – review & editing, Funding acquisition, Visualization.

## Acknowledgements

We acknowledge Dr. Julia Mannheim, Dr. Rebecca Rock, Dr. Neele Hübner, Dr. Andreas Dieterich, Ines Herbon, Stacy Huang, Sandro Aidone, and Linda Schramm for their administrative and technical support. We thank the Radiopharmacy department for tracer production. This work is also part of the Ph.D. thesis of Tudor M Ionescu.

