## supplemental information for "Neurovascular Uncoupling: Multimodal Imaging Delineates the Acute Effects of MDMA"

### 1. Supplementary Methods

**Schiffer rat brain atlas**

The rat brains were subdivided into several brain regions according to the Schiffer rat brain atlas. A total of 54 regions were selected. A list of regions, their volumes and abbreviations can be found in Supplementary Table 1.

**Supplementary Table 1: List of regions selected according to the Schiffer rat brain atlas.**

| Brain region (ROI) | Hemisphere | ROI volume [$mm^3$] | Abbreviation |
|---|---|---|---|
| Nucleus Accumbens | left | 7.9 | NAc |
| | right | | |
| Amygdala | left | 21.1 | Amyg |
| | right | | |
| Caudate Putamen | left | 43.5 | CPu |
| | right | | |
| Auditory Cortex | left | 27.5 | Au |
| | right | | |
| Cingulate Cortex | left | 14.5 | Cg |
| | right | | |
| Entorhinal Cortex | left | 59.0 | Ent |
| | right | | |
| Insular Cortex | left | 21.1 | Ins |
| | right | | |
| Medial Prefrontal Cortex | left | 6.3 | mPFC |
| | right | | |
| Motor Cortex | left | 32.6 | M1 |

| | | | |
|---|---|---|---|
| | right | | |
| Orbitofrontal Cortex | left | 18.9 | OFC |
| | right | | |
| Parietal Cortex | left | 7.6 | PaC |
| | right | | |
| Retrosplenial Cortex | left | 18.9 | RS |
| | right | | |
| Somatosensory Cortex | left | 71.6 | S1 |
| | right | | |
| Visual Cortex | left | 36.1 | V1 |
| | right | | |
| Anterodorsal Hippocampus | left | 25.1 | CA1 |
| | right | | |
| Posterior Hippocampus | left | 9.8 | CA1-p |
| | right | | |
| Hypothalamus | left | 18.4 | Hyp |
| | right | | |
| Olfactory Cortex | left | 14.0 | OC |
| | right | | |
| Superior Colliculus | left | 7.1 | SC |
| | right | | |
| Midbrain | left | 11.4 | MB |
| | right | | |

| | | | |
|---|---|---|---|
| Ventral Tegmental Area | left | 5.5 | VTA |
| | right | | |
| Cerebellum – grey matter | left | 75.0 | CG |
| | right | | |
| Cerebellum – white mater | left | 23.4 | CW |
| | right | | |
| Inferior Colliculus | left | 5.7 | IC |
| | right | | |
| Thalamus | left | 30.7 | Th |
| | right | | |
| Periaqueaductal Gray | - | 9.9 | PAG |
| Septum | - | 9.4 | Sep |

**Radiotracer synthesis**

[$^{11}$C]DASB synthesis was performed in a modified procedure [1] compared to the report by Wilson et al. [2]. In brief, a solution of 2 mg of precursor in 500 µL DMSO was used to trap [$^{11}$C]methyliodide. After the reaction was heated to 100 °C for 2 min, dilution was performed with 1.5 mL of HPLC eluent (3 mM $Na_2HPO_4$ with 64 % acetonitrile). Purification was performed on a Luna C18 column (250 mm x 10 mm, Phenomenex, Torrance, CA, USA). The isolated peak was diluted afterwards using 70 mL water with 20 mg sodium ascorbate, then loaded onto a conditioned Strata-X cartridge (Phenomenex), eluted with 0.5 mL of ethanol and diluted with 5 mL of phosphate-buffered saline.

[$^{11}$C]DASB injected activities were 175 ± 66 MBq/mL, and molar activities were 86 ± 25 GBq/µmoL at the time of injection, resulting in an average dose of 364 MBq per animal. [$^{11}$C]DASB was applied using a bolus plus constant infusion protocol with an initial bolus of 0.58 mL over the first 20 seconds and a constant infusion of 15 µL/min over the remainder of the scan ($k_{bol}$ = 37.8 min).

The $^{18}O(p,n)^{18}F$ nuclear reaction using [$^{18}$O]water (Rotem, Leipzig, Germany) was employed to prepare [$^{18}$F]fluorine as [$^{18}$F]fluoride using the PETtrace cyclotron (GE Healthcare, Uppsala, Sweden).

[$^{18}$F]FDG was synthesized in a TRACERlab $MX_{FDG}$ synthesizer (GE Healthcare, Liège, Belgium) using mannose triflate (ABX, Radeberg, Germany), as described previously [3]. Quality control after synthesis was executed in accordance with GMP guidelines.

For [$^{18}$F]FDG, a constant infusion protocol was applied, the tracer (150 MBq/mL pre-scan dose) being infused at a rate of 8 µL/min over 100 minutes, resulting in total doses of 120 MBq per animal [4].

**Temperature monitoring**

The temperature was documented in 10-minute intervals for both the MDMA cohort in a subset of 12 scans and for the PBS cohort (Supplementary Figure 1). No major changes of temperature induced by the MDMA challenge were observed.

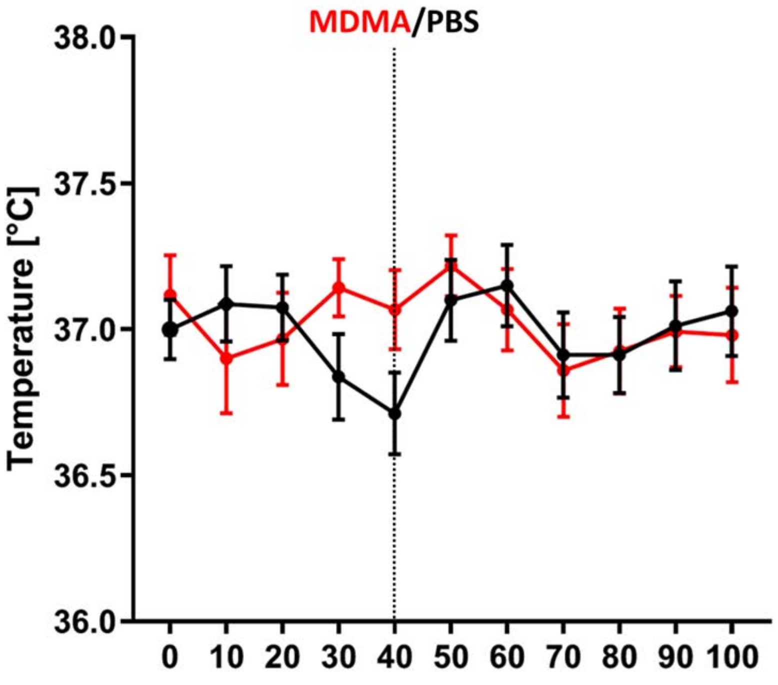


**Supplementary Figure 1: Temperatures over the course of the scan for MDMA (n = 12, red) and PBS (n = 8, black).** The data are presented as mean ± SEM.

**Preprocessing**

All fMRI and dynamic PET datasets were first realigned using SPM. Average images were generated for each fMRI and PET dataset . Using these average images and the T2-weighted anatomical references binary masks were created for extracting the brain from each fMRI, PET and anatomical scan. Subsequently, the skull-stripped fMRI and PET datasets were coregistered to the respective skull-stripped anatomical reference scans using SPM. Afterwards, the anatomical references scans were used to calculate spatial normalization parameters to the Schiffer rat brain atlas [5] which were then used to spatially normalize the fMRI and PET datasets. Finally, all fMRI and PET datasets underwent spatial smoothing using a 1.5 x 1.5 x 1.5 $mm^3$ Gaussian kernel, corresponding to the spatial resolution of the PET datasets [6].

[$^{11}$C]DASB $BP_{ND}$ values were calculated using the following formula:

$BP_{ND} = \frac{C_T - C_R}{C_R} = \frac{C_T}{C_R} - 1 = DVR - 1$, where

- $BP_{ND}$ is the non-displaceable binding potential
- $C_T$ is the concentration of the tracer in the tissue of interest
- $C_R$ is the concentration of the tracer in the reference region
- $DVR$ is the distribution volume ratio

The cerebellar gray matter was used as a reference region as previously recommended for rats [7]. For [$^{18}$F]FDG PET the cerebellum was used as a reference region, since it has been shown to outperform whole-brain normalization, which bears several caveats [8].

**Subject-level GLM PET analysis**

In addition to the voxel-wise analysis generated from static PET images presented in the main manuscript, an analysis was performed using the dynamic datasets. The analysis was performed similarly to the fMRI dataset by generating first-level contrasts between the baseline (30-40 minutes after scan start) and the 10-minute periods after MDMA administration. The first-level contrasts were then used to produce group-level contrasts in a second-level analysis.

**PBS cohort**

To generate control datasets, simultaneous PET/fMRI scans were acquired using the same protocols using both $[^{11}C]$DASB (n = 4) and $[^{18}F]$FDG (n = 4). One dataset from each cohort was excluded due to technical issues. The preprocessing and analysis of the remaining scans (n=3 per group) was performed identically to the MDMA cohort.

#### 2. Supplementary Results

**Voxel-wise PET maps**

Voxel-wise PET maps for $[^{11}C]$DASB indicating $BP_{ND}$ values and for normalized $[^{18}F]$FDG uptake were generated for the baseline (30-40 minutes after scan start) and for each 10-minute period after MDMA administration. The respective $[^{18}F]$FDG PET uptake maps can be seen in Supplementary Figure 2. The respective $[^{11}C]$DASB PET $BP_{ND}$ maps are shown in Supplementary Figure 3. The figures illustrate increase in $[^{18}F]$FDG uptake and the gradual decrease in $[^{11}C]$DASB binding elicited by the acute MDMA challenge.

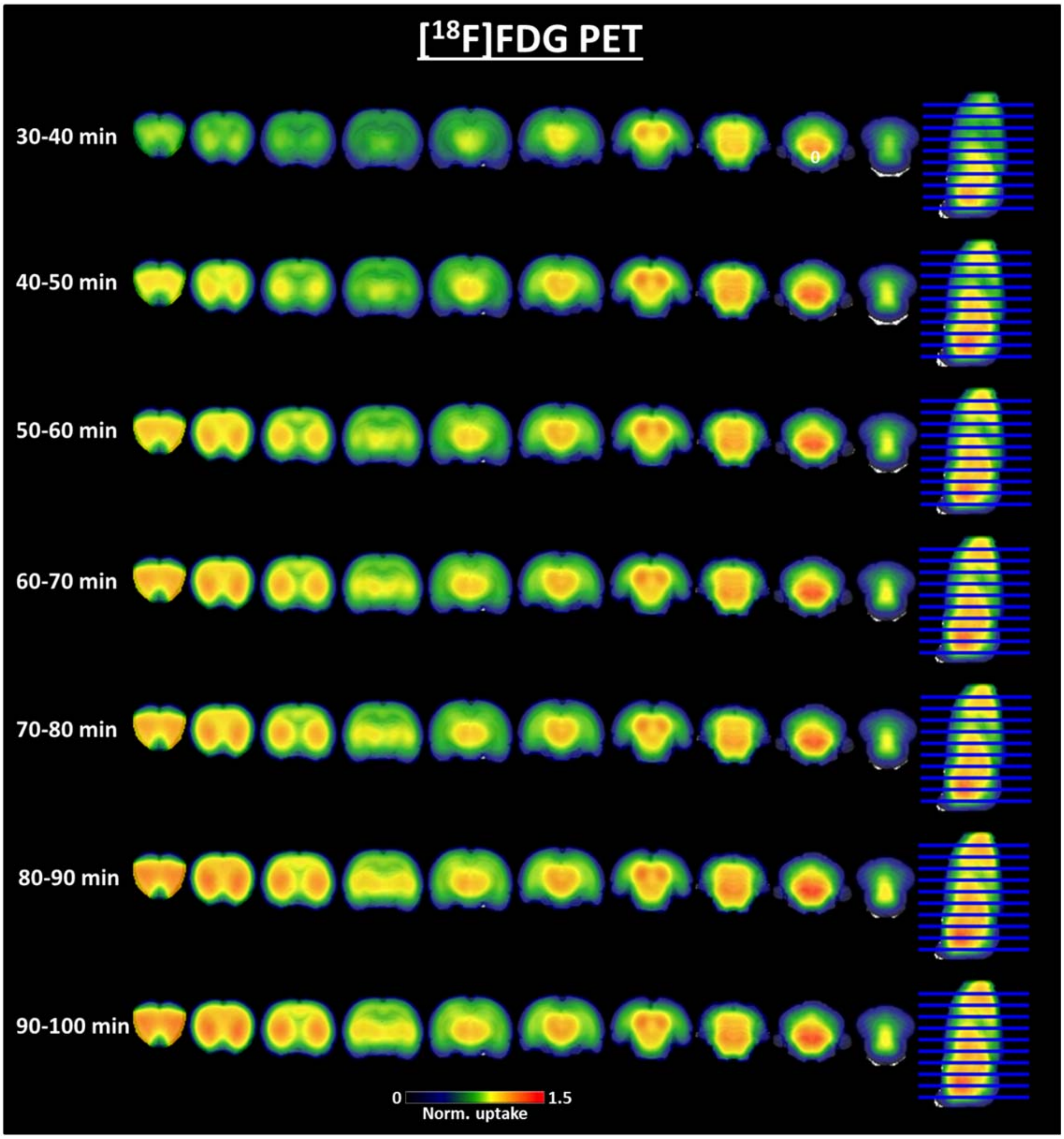


**Supplementary Figure 2: Group-average normalized [$^{18}$F]FDGuptake maps for each analyzed time period.** The uptake maps were calculated voxel-wise by normalizing the value of each voxel through the average uptake in the cerebellum. The maps indicate the gradual increase in glucose metabolism after the MDMA challenge, particularly in anterior cortical and subcortical areas.

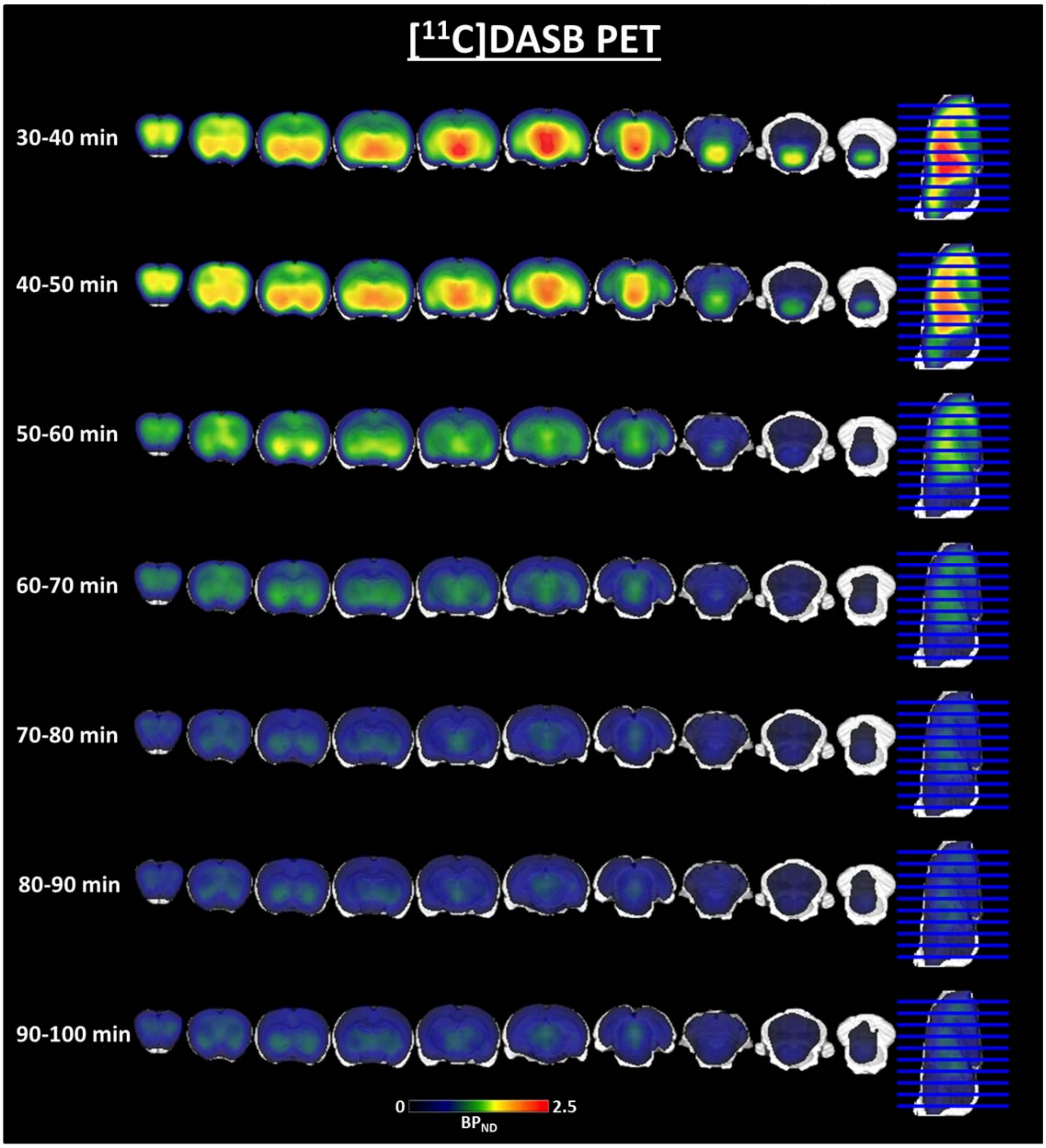


**Supplementary Figure 3: Group-average [$^{11}$C]DASB binding maps for each analyzed time period.** Voxel-wise $BP_{ND}$ values were calculated as DVR-1 using the cerebellar gray matter as a reference region. The images indicate the decreased SERT availability after acute MDMA injection.

#### PBS cohort

For group-level analysis, we merged the fMRI datasets acquired with [$^{11}$C]DASB and [$^{18}$F]FDG, therefore a total of 6 scans were used.

Supplementary Figure 4 indicates average BOLD time-courses extracted from cerebral and extracerebral areas and plotted for comparison to the respective time-courses extracted from the MDMA cohort. Additionally, two-sample t-tests were performed comparing the alterations of the BOLD signal to baseline between both cohorts. Significantly higher decreases were observed for the MDMA cohort across the areas indicated in the main manuscript.

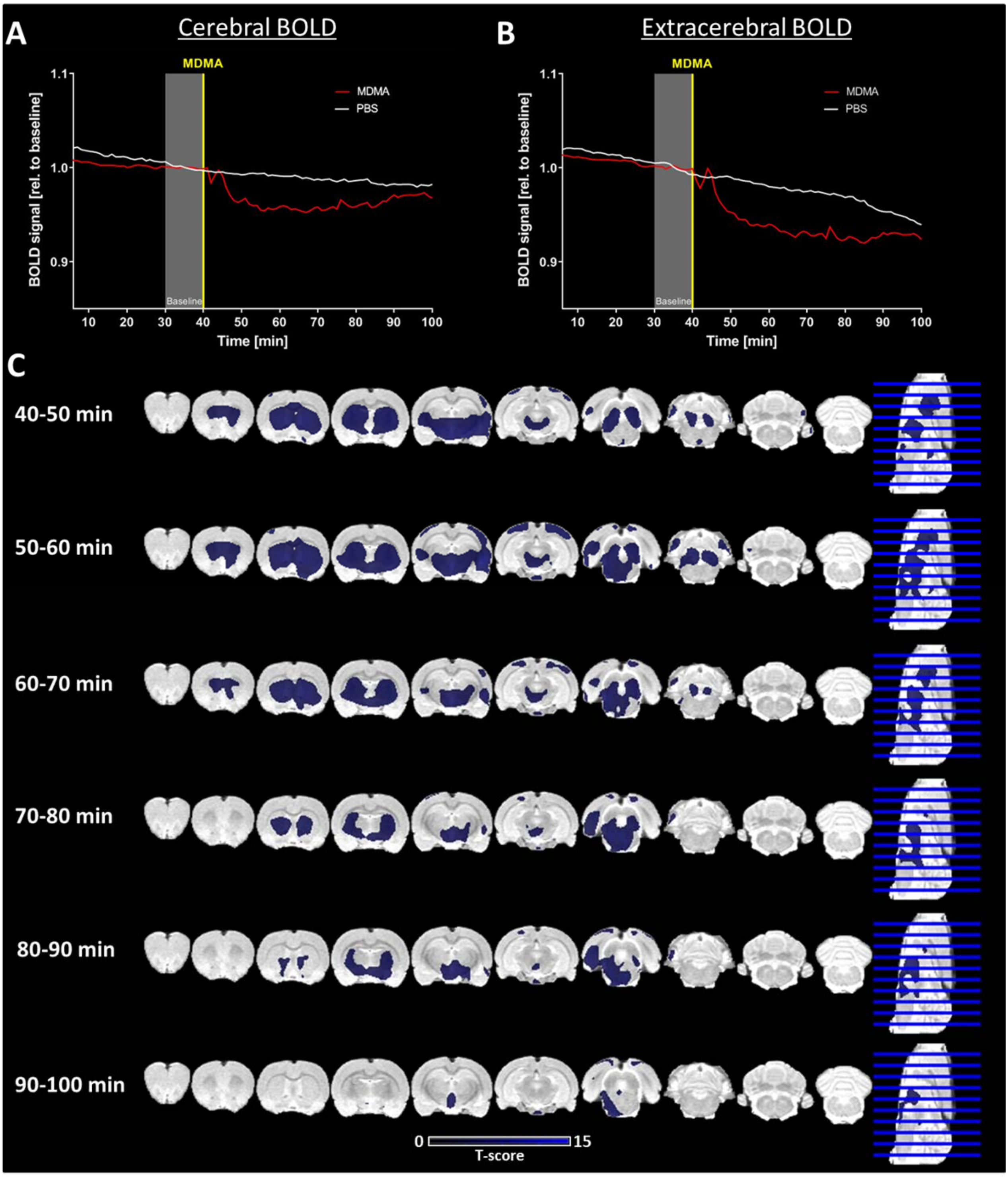


**Supplementary Figure 4: Comparison between BOLD-fMRI signal changes in MDMA and PBS cohorts. (A)** Group-average whole-brain BOLD signal over the courses of the scans normalized to baseline from minute 30 to 40. **(B)** Group-average extracerebral BOLD signal over the courses of the scans normalized to baseline from minute 30 to 40. **(C)** Voxel-wise two-sample t-tests between changes to baseline (30-40 minutes after scan start) at each subsequent time-period for MDMA and PBS cohorts (MDMA > PBS, $p < 0.001$ (voxel-level), $p < 0.05$ FWE cluster-level correction).

Linear temporal decreases, potentially driven by longitudinal effects of the used anesthesia, were observed in both cohorts (Supplementary Figure 4A and B). In contrast to the strong decreases induced by MDMA, however, no alterations were observed after the PBS challenge, as confirmed by the between-sample voxel-wise analysis (Supplementary Figure 4C).

Group-average regional [$^{18}$F]FDG normalized uptake and [$^{11}$C]DASB $BP_{ND}$ values were generated for the PBS cohorts. Neither readout indicated alterations induced by the PBS injection (Supplementary Figure 5).

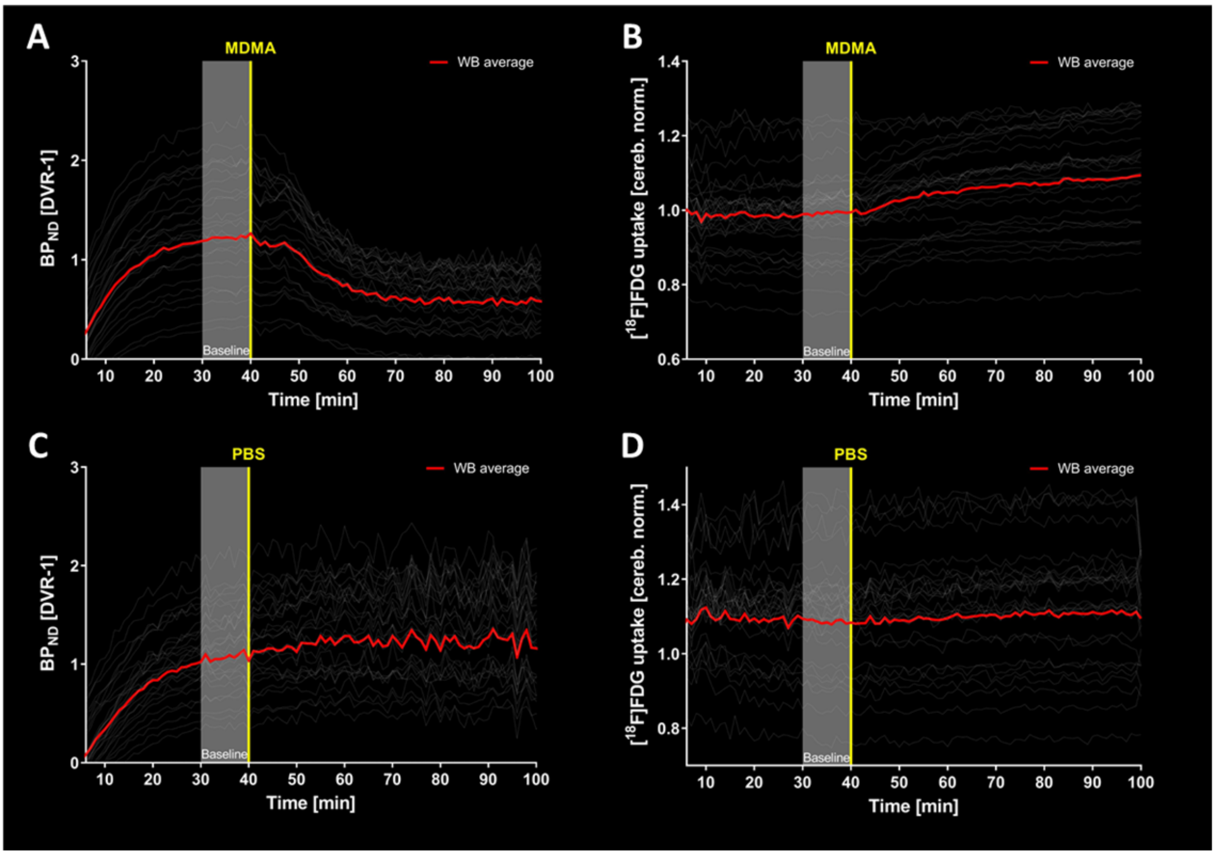


**Supplementary Figure 5: Comparison between MDMA and PBS effects on [$^{11}$C]DASB binding and [$^{18}$F]FDG uptake. (A)** Group-average [$^{11}$C]DASB binding potentials for all regions and whole brain in MDMA cohort. **(B)** Group-average [$^{18}$F]FDG uptakes for all regions and whole brain in MDMA cohort. **(C)** Group-average [$^{11}$C]DASB binding potentials for all regions and whole brain in PBS cohort. **(D)** Group-average [$^{18}$F]FDG uptakes for all regions and whole brain in PBS cohort. Sample sizes: [$^{11}$C]DASB MDMA: n = 11; [$^{18}$F]FDG MDMA: n = 15; [$^{11}$C]DASB PBS: n = 3; [$^{18}$F]FDG PBS: n = 3.

**Subject-level GLM PET analysis**

Three single-subject first-level GLM analysis results and the second-level group-average results are shown in Supplementary Figure 6 and Supplementary Figure 7, respectively, indicating decreases in [$^{11}$C]DASB binding potentials and [$^{18}$F]FDG uptakes 50-60 minutes following MDMA administration compared to baseline.

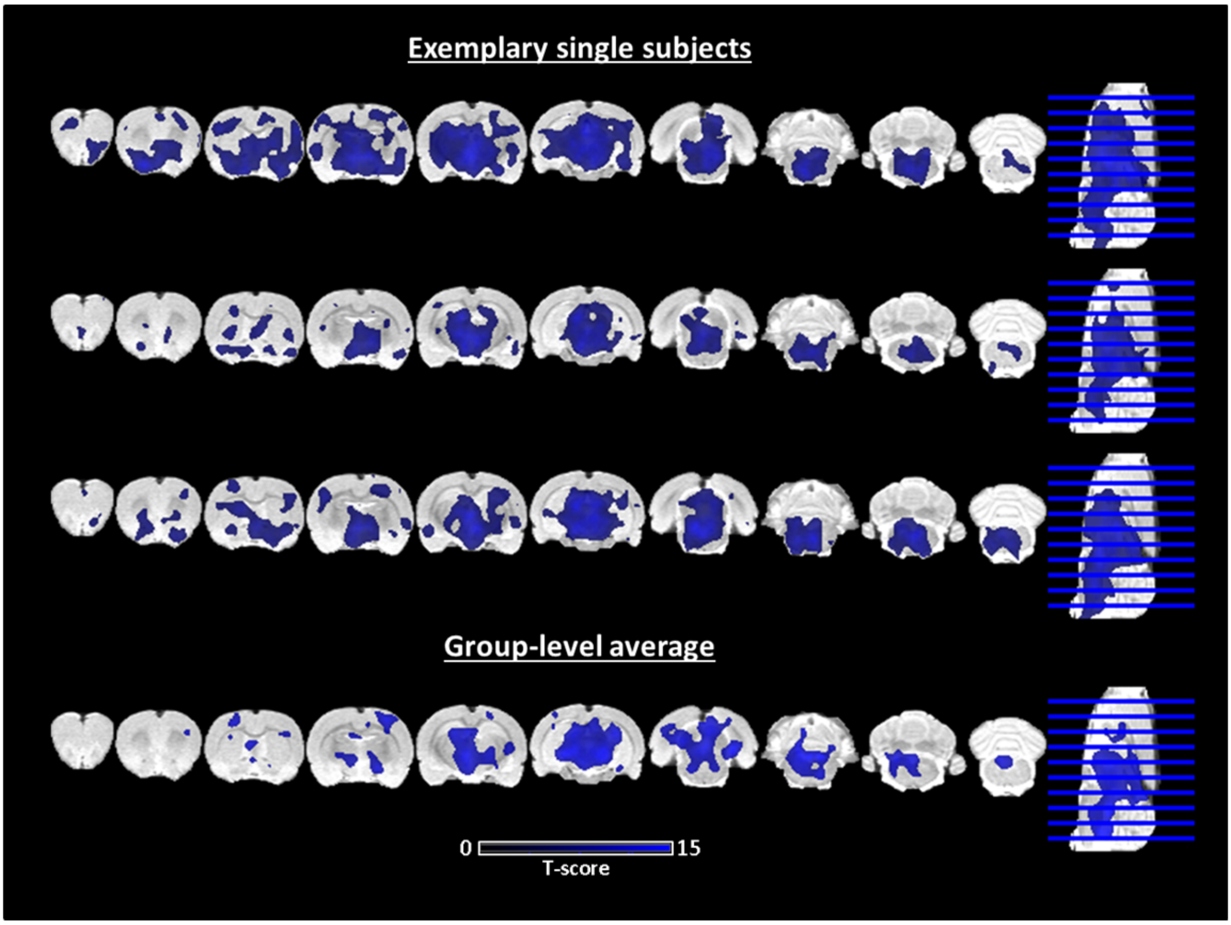


**Supplementary Figure 6: First-level analysis of [$^{11}$C]DASB PET datasets indicating decreased binding potentials in three exemplary subjects and in the generated group-average readout in the period 50-60 minutes after the MDMA challenge compared to baseline ($p < 0.05$, FWE corrected).**

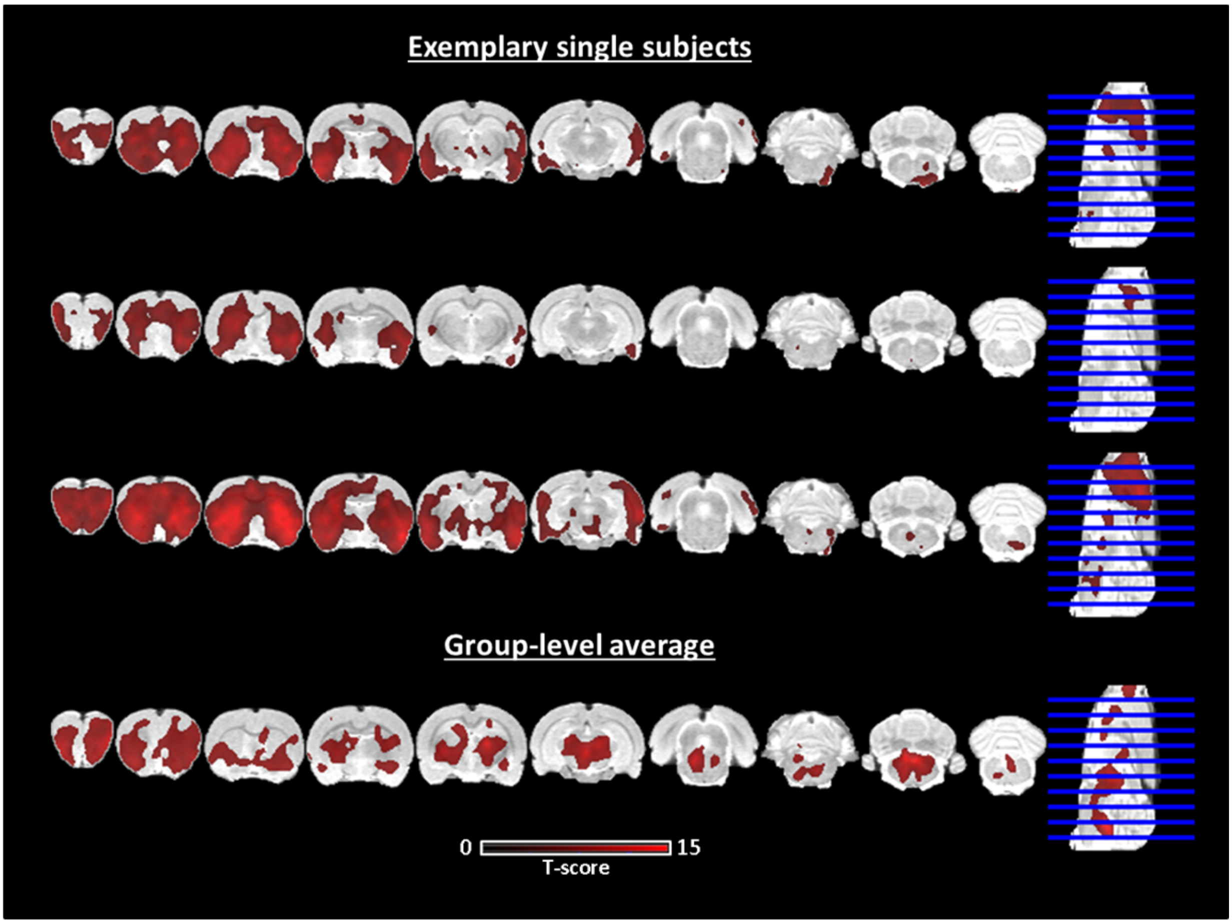


**Supplementary Figure 7: First-level analysis of [$^{18}$F]FDG PET datasets indicating increased uptakes in three exemplary subjects and in the generated group-average readout in the period 50-60 minutes after the MDMA challenge compared to baseline ($p < 0.05$, FWE corrected).**

### Supplementary Discussion

#### The power of multimodal imaging

We show the ability of multimodal imaging to elucidate *in vivo* effects of drugs due to the lack of reliance on hemodynamics. While BOLD-fMRI can be a very useful method and exhibits various advantages to PET in terms of spatial or temporal resolution, in addition to not requiring the administration of radioactive substances, its convoluted substrate can make its mapping onto neuronal activity challenging, as demonstrated by our study. The inclusion of PET in the equation offers a variety of benefits, as previous work has also shown. When[$^{18}$F]FDG fPET and fMRI were used simultaneously for activation studies, different activation patterns were observed [6]. For fMRI-derived resting-state functional connectivity, PET can offer crucial additional information on neurotransmitter modulation of specific networks [9, 10] and can be used to derive complementary networks on other physiological levels [4, 11]. In terms of pharmacological research, PET/fMRI imaging has yielded important complementary findings in several previous studies [12].

Psychiatric conditions are an increasing burden in modern societies [13, 14]; therefore, research on novel therapies and drugs to treat these conditions is required. Imaging has helped provide promising biomarkers for psychiatric drug research in recent years and has contributed to the emergence of psychedelic substances as plausible candidates for novel therapeutic strategies in psychiatry [15]. The present study demonstrates that advanced imaging technology can offer tremendous insight in understanding the mechanism of drugs compared to unimodal scans. Future research employing similar study protocols in either clinical or preclinical pathological cohorts would be of significant interest to elucidate how the acute effects of drugs such MDMA revealed by PET in addition to fMRI correlate with therapeutic outcomes. Importantly, using the current study design employing constant tracer infusion, effects can also be reliably mapped in single patients, as opposed to the readout solely on group-level when applying the tracer as a single bolus, thus ensuring the clinical translatability. While not in the scope of the study, correlating the effects of the different readouts between subjects may offer further important insights.

**Limitations**

The present study was performed on anesthetized animals. Preclinical studies offer many advantages compared to human studies, such as high cohort homogeneity regarding age, strain, and gender. However, the effects of the used anesthesia must be considered especially when compared to similar human awake studies. Generally, interactions between the drug and the used anesthesia cannot be excluded [16]. Studies comparing the readout of the MDMA challenge under awake conditions and different anesthesia protocols are therefore necessary for the future. Nonetheless, we kept the anesthesia at levels indicated previously for fMRI scans under isoflurane in terms of physiology and readout [17, 18]. The patterns of the BOLD decreases were in concordance with previous data acquired in awake humans [19], as described above.

Furthermore, our BOLD scans did not have whole-brain coverage; therefore, information from parts of the cerebellum was missed. Additionally, using a surface coil likely led to a poorer BOLD signal quality in deeper areas than the cortex. Studies covering all brain areas with volume coils may reveal additional information on the effects of this drug on the hemodynamic level.
